# A phylogenetic analysis of MCTP proteins: from amino acid sequence to function

**DOI:** 10.1101/2020.01.13.904789

**Authors:** José Luis Téllez Arreola, Argel Estrada-Mondragón, Ataúlfo Martínez Torres

## Abstract

MCTPs (Multiple C2 domain proteins with two transmembrane regions) are evolutionarily and structurally related to other C2 proteins which play fundamental roles in exocytosis and membrane trafficking, however their specific role has been little studied. This work points out possible functional implications of MCTPs by comparing their primary amino acid sequence and functional domains. MCTP amino acid sequences were identified in non-chordates and chordates. The primary sequences grouped in three classes: MCTP, MCTP-1 and MCTP-2. MCTP is present only in non-chordates, while MCTP-1 and MCTP-2 are present in chordates. MCTP genes emerged early in metazoan evolution and are well conserved across species including humans. Genomic analysis of diverse species of representative phyla showed that the three C2 domains (C2A-C2C) and transmembrane regions (TMR) are well conserved. The C2 domains have eight β strands as well as aspartate residues known to bind calcium. Interestingly, we identified a lysine-rich cluster, also known as polybasic cluster in C2A and C2B, which is known to bind lipids in other proteins. We also describe the phylogenetic distribution of MCTPs and analyze conserved domains and their predicted secondary structure in metazoans. We highlight important motifs that have not been previously described in MCTPs C2A and C2B domains that suggest MCTPs potentially bind phospholipids. Our observations show MCTPs are proteins widely distributed in eukaryotic organisms and may play an important role in membrane fusion or exocytosis.

## Introduction

Cellular signaling through proteins that possess calcium-binding domains is fundamental for multiple processes such as exocytosis, vesicular trafficking, protein kinase and phospholipase activities, among many others. The C2 domain is present in many proteins involved in synaptic transmission such as synaptotagmins (Bagur and Hajnóczky, 2017; Evans et al., 2016; Nalefski and Falke, 1996; Shin et al., 2005). This domain was originally described as the second of four calcium-binding domains of the calcium-dependent protein kinase C (Nalefski and Falke, 1996); however, it is now well known that this domain is widespread in diverse eukaryotic proteins that could act or not as calcium sensors, triggering diverse cellular responses (Bagur and Hajnóczky, 2017). Molecular and functional studies have revealed the role of proteins that possess C2 domains in different signaling cascades; nevertheless, multiple questions remain. C2 proteins involved in membrane trafficking, such as synaptotagmins and ferlins, include a transmembrane region and at least two C2 domains whose calcium affinity is higher in the presence of phospholipids (Rizo and Sudhof, 1998). A novel family of C2 proteins (MCTPs) was discovered by *in silico* genomic sequence analyses (Shin et al., 2005). They present three conserved C2 domains (C2A, C2B and C2C) facing the cytoplasm, one- or two-transmembrane regions that anchor the protein to intracellular vesicles and one short C-terminus sequence. Vertebrates have two MCTP coding genes (*MCTP1* and *MCTP2*). While, invertebrates such as *Drosophila melanogaster* and *Caenorhabditis elegans* have only one MCTP gene (Shin et al., 2005). The cellular distribution of MCTPs remains little known. They appear to be located in intracellular vesicles and endoplasmic reticulum, which are a suitable positions for sensing changes in calcium concentration and to play a role in vesicular trafficking (Shin et al., 2005), yet their functions are not known.

Assessment with selective antibodies for each isoform proved that MCTPs are widely distributed in vertebrates, especially in striated muscle and heart (Qiu et al., 2015; Shin et al., 2005). This is interesting since a 2.2 Mb deletion in the 15q26.2 chromosome, in which the *MCTP*2 gene is located, causes coarctation of the aorta and hypoplastic heart syndrome (Lalani et al., 2013). Furthermore, the importance of the *MCTP*2 gene for proper cardiogenesis in frogs, was proved by knocking down the expression of the gene with morpholinos, which yielded embryos with a heart condition resembling coarctation of the aorta (Lalani et al., 2013). In other studies, single nucleotide polymorphisms in the *MCTP2* gene were associated with schizophrenia and bipolar disorders, yet its role in evoking these psychiatric diseases has been elusive (Djurovic et al., 2009). In 2015, a study described the distribution of the MCTP-1 in rats. It showed that MCTP-1 is widely distributed in the brain and is found in secretory vesicles and endosomes of neurons (Qiu et al., 2015). More recently, it was found that human MCTP2 transiently expressed in COS7 cells is located in the endoplasmic reticulum and peroxisomes (Joshi et al., 2018).

Studies of the role of MCTPs have also been carried out in invertebrates. Only one gene has been identified in the genome of *D. melanogaster*, and a gene-trap study showed that it is expressed in the accessory cells of the olfactory organ. A P-transposon insertion at the 3’end of the non-coding region of the MCTP gene induces high lethality in the fly (Tunstall et al., 2012). In contrast, other allelic mutants of the MCTP gene are viable (Genç et al., 2017) and this evidence allowed the authors to suggest that this protein is necessary for presynaptic homeostatic plasticity and to set off the baseline for neurotransmitter release. This study also established that MCTP is located in the endoplasmic reticulum of motoneurons, where it senses changes in calcium concentrations, thus facilitating neurotransmitter release. But, further evidence is required to conciliate these contrasting evidences.

In *C. elegans* the study of MCTP is more limited. Two isoforms are predicted to be expressed by the use of two alternative promoters, but the gene expression pattern or cellular localization of the protein has not been determined. Nevertheless, a massive antisense RNAi screening disclosed that when expression of MCTP is knocked down (Maeda et al., 2001), larval lethality is high, consistent with the study of Tunstall et al. (2012) in *D. melanogaster* and suggesting the relevance of this gene for proper development.

Much has been learned about the structure and role of proteins that include the C2 domain, originally identified in the PKC proteins (Nishizuka, 1988). The structure of this domain was revealed by high-resolution X-ray diffraction of synaptotagmin crystals, a family of proteins involved in synaptic transmission (Alwarawrah et al., 2017; Corbalan-Garcia and Gómez-Fernández, 2014; Evans et al., 2016; Sutton et al., 1995). C2s are conformed by approximately 130 amino acids and include aspartate residues that coordinate calcium binding. MCTPs are structurally related to synaptotagmins, extended-synaptotagmins (Esyts) and ferlins (Min et al., 2007), proteins that possess two or six C2 domains in tandem, respectively. Nevertheless, the function of the C2 domains in the MCTPs as well as the functional role of these proteins are not fully understood.

In this study, we present a phylogenetic analysis of the MCTP protein family, aiming to shed light on the structural characteristics of this emerging family of calcium sensors.

## Methods

### Identification of the MCTP protein sequences

All the sequences for MCTP proteins were obtained by using SMART (Schultz et al., 1998) and applying the *C. elegans* MCTP protein sequence (**<u>NP491908.2</u>**) as template. The presence of three C2 domains in their primary sequence and the predicted transmembrane regions were the main criteria to identify MCTP sequences in other species.

### Multiple sequence alignment

Sequence alignments were performed in COBALT using default parameters. COBALT uses multiple alignment of sequences, a conserved domain database, protein motif databases and local sequence similarity using RPS-BLAST, BLASTP and PHI-BLAST (Papadopoulos and R., 2007). Jalview was used to color sequence alignments derived from COBALT results. Zappo color schemes were used to classify amino acids according to their physicochemical properties (Waterhouse et al., 2009).

### MCTP phylogenetic analysis

We used multiple sequences obtained by COBALT as input to generate phylogenetic trees, to analyze the spread of MCTP proteins in all different species. The phylogenetic tree of experimental model organisms and C2 domains tree were generated by using muscle alignment program, maximum likelihood method and Bayesian inference (M.Bayes3.2) or bootstrapping (100 replicates) within phylogeny.fr platform as needed.

### MCTP secondary structure prediction and structural modeling

We used PSIPRED software for secondary structure prediction, and RaptorX Structure Prediction software for modelling the tertiary structure of MCTP from *C. elegans* (**NP491908.2**) using the default parameters of the server (Jones, 1999; Källberg et al., 2012). The automatically generated pdb model was manually cropped in Molsoft ICM-Pro v3.8-3 keeping the integrity of all the predicted domains in a common configuration. One calcium molecule was loaded from ChemSpider into the ICM project (CSID: 4573905), and fitted in the MCTP C2A domain using as guide the structure of the C2A domain of Synaptotagmin-7 (PDB: 6ANJ, Voleti et al., 2017) using the function Pocket Finder of ICM (Dey and Chen, 2011). The final model was submitted to Yasara energy minimization server (Krieger et al., 2009) and the images generated in PyMOL v2.2.2 Schrodinger LLC (Schrödinger, San Diego, CA).

## Results

### MCTPs are found in metazoan but not in unicellular eukaryotes

MCTP protein sequences were found to be widespread in metazoans, from nematodes to humans. We found no evidence of MCTPs in prokaryotic organisms or unicellular eukaryotes. We generated an evolutionary tree of MCTPs from different organisms that are used as experimental biological models including the sea squirt *Ciona intestinalis,* a tunicate that represents the base line of chordates (Fig. 1). The tree analysis shows the divergence of the MCTP protein family into three major groups, MCTPs class1 and class 2 are present in chordates such as bony fish, reptiles, amphibians, birds and mammals (rodents, ungulates and primates including humans, Figure 1, and supplementary list). The clas MCTP was found only in invertebrates (nematodes, flies and mollusks) including parasitic and free-living nematodes.

**Fig. 1.**
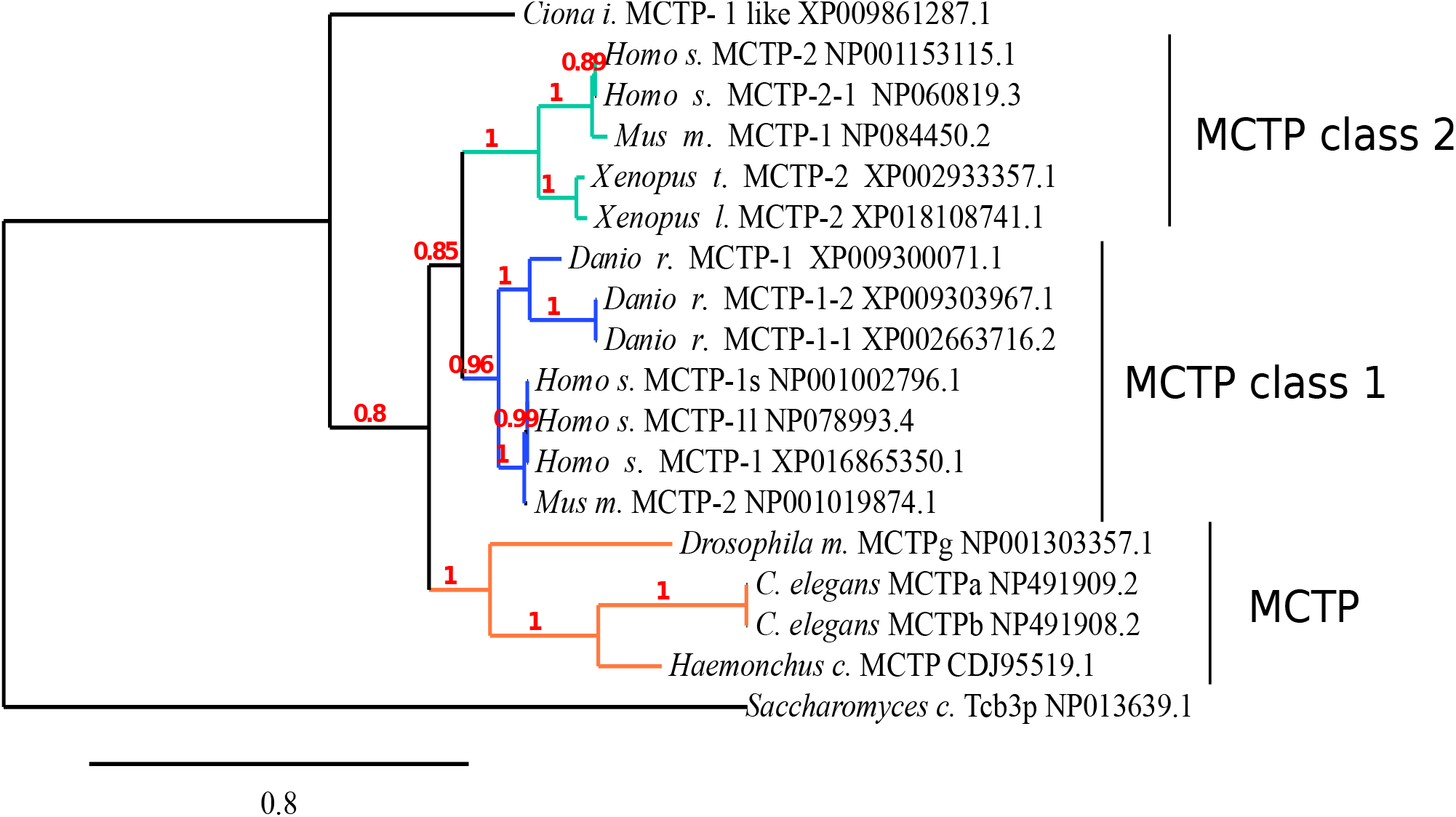
Phylogenetic analysis of the MCTPs proteins sequences. All sequences selected from the NCBI appear with their accession number. The tree was generated using maximum likelihood method and Bayesian inference using 100 bootstrap replicates, only the model species tree is shown because the patter is the same than the General analysis. Sequence of *C. intestinalis* was used as outgroup. The scale bar represents the length distance units of the tree branches.

### Analysis and modeling of the conserved functional domains of the MCTPs

As previously mentioned, MCTP proteins possess three cytosolic C2 domains in tandem and two transmembrane regions (Fig. 3A). We aligned the amino acid sequences corresponding to individual C2 domains (C2A, C2B and C2C) from the different three classes of MCTPs as well as their C-terminus to determine the degree of conservation of the amino acid sequence of the transmembrane regions (Fig 3B-G). A comparison of the C2 domains revealed a high conservation in the primary sequence, although the degree of conservation is different among them. In general, the C2C domain is the most conserved, followed by C2A and finally C2B; however, calcium binding sites are more conserved in C2A than in C2C (Fig. 3 B-D). The number and position of the aspartate residues that potentially bind calcium in the MCTPs was analyzed and compared to the C2A of Synaptotagmin 1 (**P21579.1**), and we found that the position of five of these residues are partially conserved in the three C2 domains of MCTP. For example, the aspartate residues 1 to 4 are conserved in the C2C domain, whereas the fifth presents the substitution D → E in *C. elegans, H. contortus, D. melanogaster* and *C. intestinalis* (Fig 3D); furthermore C2B exhibits only four aspartate residues instead of five and all of them show diverse amino acid substitutions, including D → A, E, H, N or T (Fig. 3C). Other MCTP2-C2Bs from *C. elegans, haemonchus contortuos, Xenopus leevis, Xenopus tropicalis* and some MCTP2 isoforms in human and mice exhibit only one or two aspartate residues, thus suggesting that they may not bind calcium (Fig. 3B-D). Amino acid residues predicted to bind calcium are well conserved in C2A and C2C in all analyzed MCTPs analyzed but remarkably that is not the case for C2B.

In C2A and C2B domains of the MCTPs, a lysine-rich domain previously named the polybasic cluster was detected (Corbalan-Garcia and Gómez-Fernández, 2014; Di Paolo and De Camilli, 2006). This domain is predicted by the presence of positive charged (K, K, K) and aromatic (Y, W) residues (Figs. 3 F-G, 4). The polybasic cluster is responsible for binding phosphatidylserine and PtdIns(4,5)P2 (PIP_2_) in calcium-binding domains of C2 proteins such as Syt1 (Corbalan-Garcia and Gómez-Fernández, 2014).

This presence of lysine-rich domain in the C2A and C2B of MCTP has opened the door for a working hypothesis to determine whether phospholipids could be necessary for calcium binding, which contrasts with the initial observations that suggested that MCTP function is independent of lipid interactions (Shin et al., 2005). Altogether, we found at least two potential PIP_2_ binding sites that resemble other C2 domains known to penetrate the plasma membrane, these PIP2 binding sites, in coordination with the aspartates present in C2A, C2B and C2C can potentially stabilize a calcium molecule (Fig. 3A-D). On the other hand, the SMART software yielded a model that suggests that the C2 domains of MCTP spans around 125 residues, whereas PSIPRED suggests that the secondary structural of the C2 domains are eight *β*-strands, that is a characteristic of the C2 domain conformation Class 2 (topology-II) in which the N and C terminus are near to the bottom (Figs. 3 A, 4).

A second criteria to establish that a protein belongs to MCTPs was the presence of two stretches of hydrophobic residues that form the transmembrane region (TMR) that allows the protein to anchor to cell vesicles and/or the plasma membrane. It has been shown that only one segment suffices to anchor the protein to intracellular vesicles (Shin et al 2005), although is unknown whether missing one of the TMRs misdirects the trafficking of the protein. The amino acid sequences of the TMRs of MCTPs were analyzed by SMART. MCTP-2 includes two TMRs located towards the C-terminus, with exception of the isoform 2 from *H. sapiens* MCTP-2, which lacks one TMR (Fig. 3E). The amino acid sequence of these TMRs is not related to other known C2 proteins involved in membrane trafficking that also include in their structure TMRs, such as synaptotagmins. In MCTPs, the second TMR is well conserved in different phyla, whereas the sequence of the first TMR is dispersed. Functional analysis using GFP-tagged versions of mammalian MCTPs previously revealed that both TMRs are functional in transfected cells (Shin et al., 2005) and direct the protein to intracellular vesicles, suggesting that one TMR suffices to anchor the protein in vesicles.

### Divergence of the C2 domain

We explored whether the amino acid sequence of C2 domains of MCTPs vary among phyla. Alignments of all three C2 domains from representative species of each clade were used to generate maximum likelihood trees. We observed three general clades: the first included the C2C domain from all species (*C. elegans*, *H. contourtus*, *C. intestinalis, X. leavis, X. tropicalis, D. rerio, M. musculus* and *H. sapiens*). The second and major clade includes C2A and C2B, thus suggesting that they may have similar functions, at least as suggested by their primary amino acid sequence. Interestingly, both C2As and C2Bs possess the putative lipid binding residues included within the lysine-rich domain disclosed by our analysis, in sharp contrast, C2Cs do not have this domain, pointing out to a structural feature that suggests different functions (Fig. 2).

**Fig. 2.**
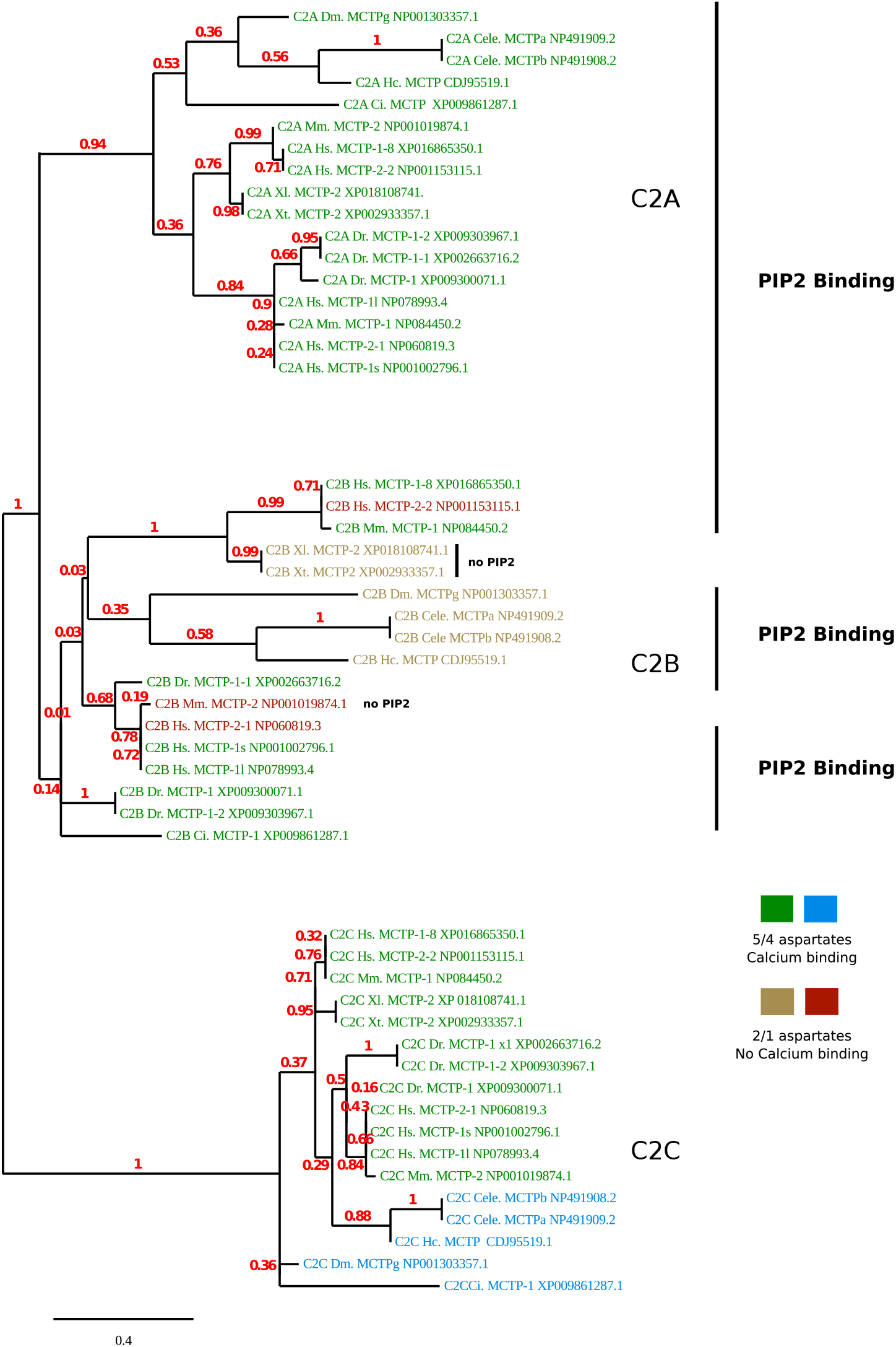
MCTP C2 domains phylogeny. A Maximum likelihood tree of C2 Domains sequences from different animal models is shown. Bootstrap inference was used making 100 replicates, values along branches are indicated in red. Putative calcium binding and Lipid binding domains were colored in the figure and indicated respectively. The scale bar represents the length of the branches of the tree.

**Fig. 3.**
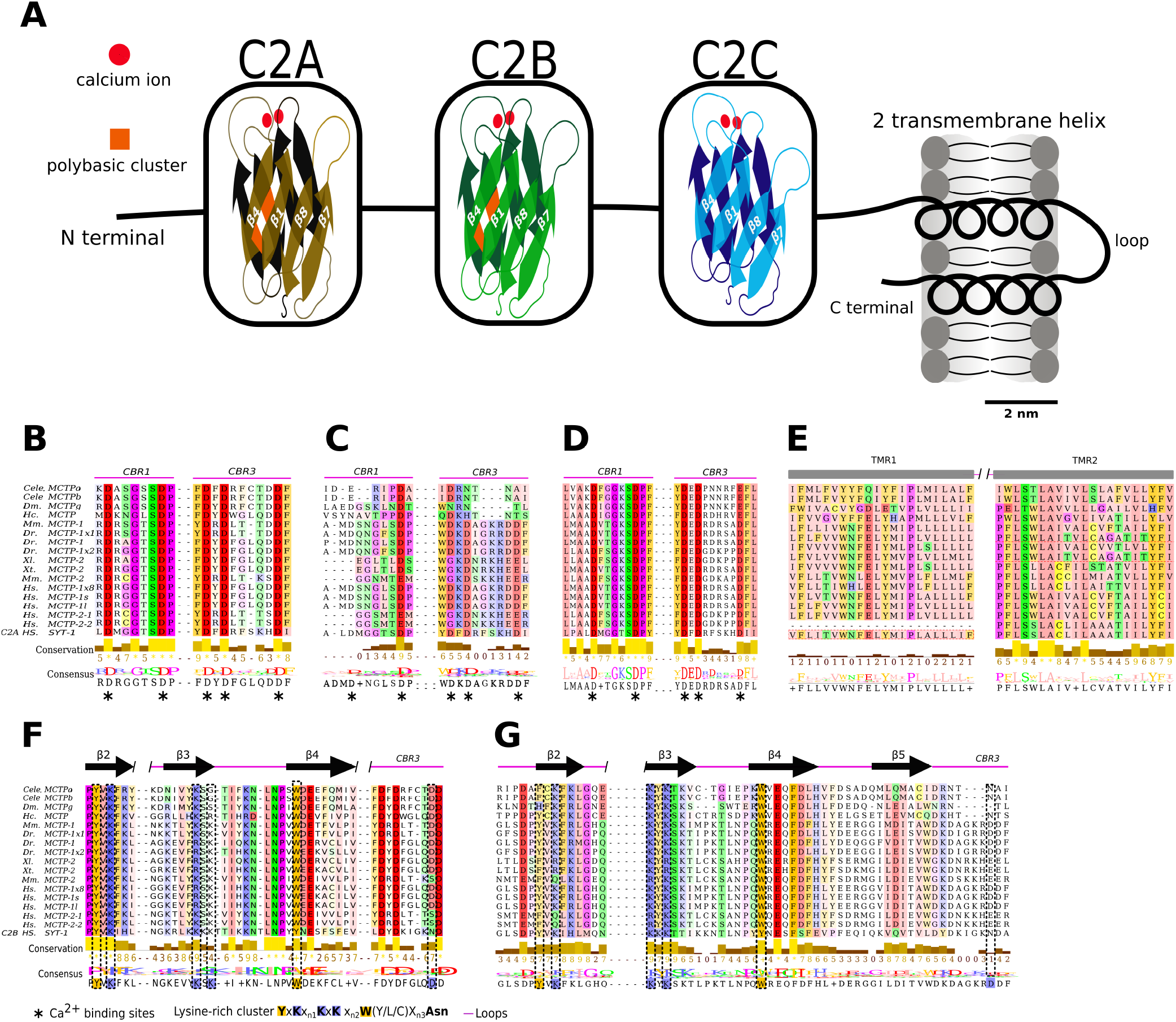
Functional Motifs in MCTPs Proteins. **A)** diagram of MCTP using consensus sequence generated by alignment of model species. Two transmembrane helix at the C- terminus anchored protein to the membrane, followed by three C2 domains. Putative calcium and lipid binding sites are shown in the scheme in C2A to C2C and C2A and C2B respectively. To determine the conservation of calcium sites **B-D)** and lipid binding pockets **F,G)** sequences from C2A and C2B from the human synaptotagmin-1 were used as a template respectively. The secondary structure was calculated using PSIPRED software. Columns marked with black asterisks are calcium binding sites in all figures, colored rectangles at the bottom of each alignment highlighted the *β*-groove motif. **E)** Alignment from TMR-1 and TMR-2 from representative species. Alignment were made using muscle and Jalview for colored them. Color scheme was zappo style. The conserved regions were calculated using SMART. Columns into gray rectangles represent the TMR regions 1 and 2, respectively.

**Fig 4.**
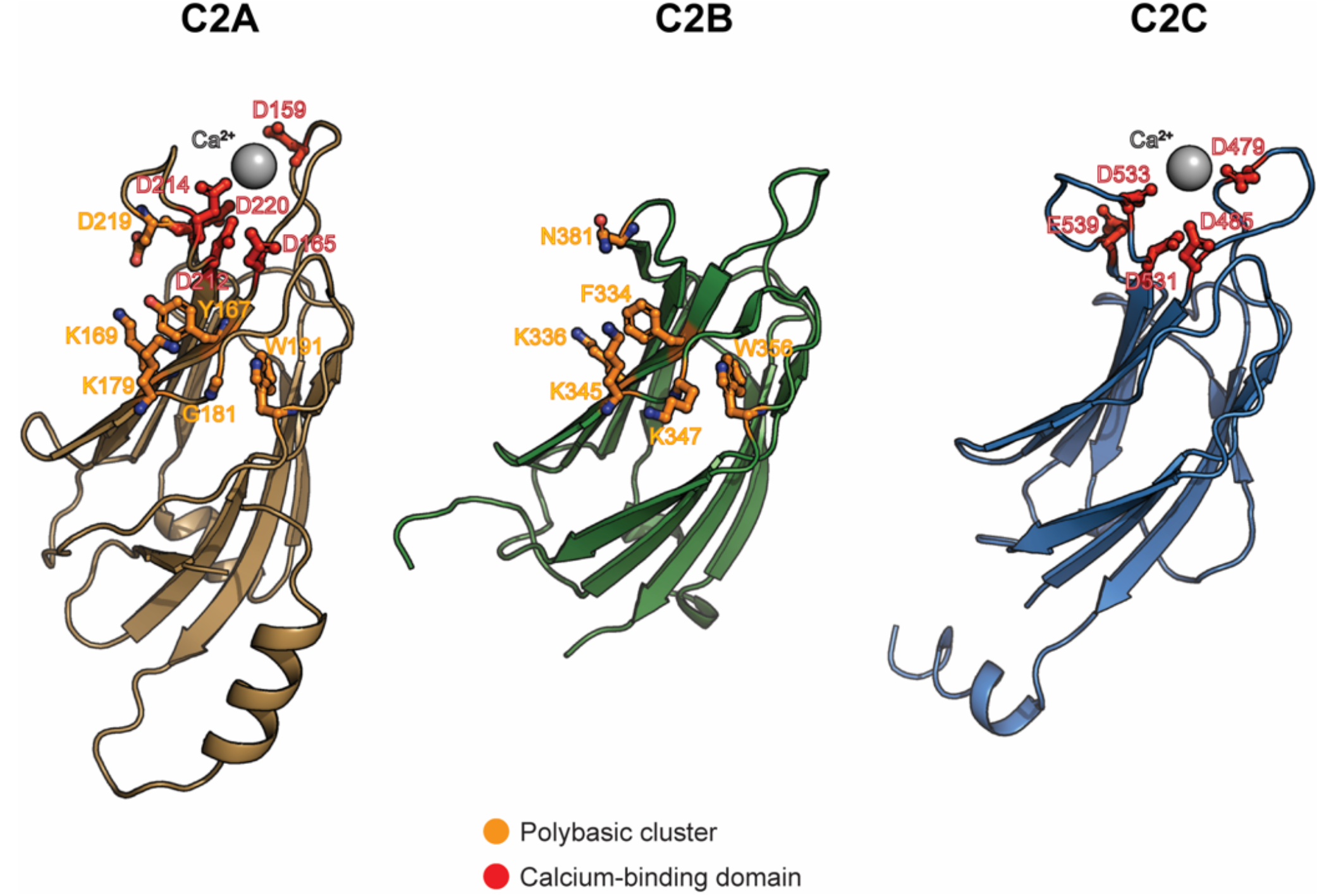
Structural model of C2 domains on MCTP. Tridimensional representation of C2A (brown), C2B (green), and C2C (blue), domains of MCTP from *C. elegans*. Calcium molecules (grey spheres) interacting with Calcium-binding domain-related residues (red) are indicated. The predicted polybasic cluster-related residues (orange) are also illustrated.

## Discussion

In the present study we provide an evolutionary analysis of the MCTP proteins. By definition, MCTPs are composed of a variable N-terminal region, three C2 domains, one or two TMRs and one short C-terminal sequence (Shin et al., 2005). We used this approach to determine peculiarities in the amino acid sequence of the MCTPs to formulate a working hypothesis to try to understand their functional role. In plants sixteen like-MCTPs classes were identified, however most of them have more than three C2 domains and three to four TMRs (Liu et al., 2017). These features are more related to other C2 proteins such as E-syts and tricalbins (Fernandez-Busnadiego et al., 2015; Saheki and De Camilli, 2017), thus, these proteins were not included in our analysis. In this work, we suggest the existence of three classes of MCTPs: MCTP, MCTP-1 and MCTP-2, that emerged from a common ancestor. Their phylogenetic distribution is disperse but different. MCTP-1 and MCTP-2 are present only in vertebrates, (bony fish, snakes, birds, rodents, ungulates and primates), whereas MCTP was observed only in invertebrates (Fig. 1). The calcium sensor proteins synaptotagmins are central molecular components of the neurotransmitter exocytotic machinery in the presynaptic terminal. They appeared early in the evolution of eukaryotes, and evidence shows that they are present even in primitive forms of eukaryotic multicellular organisms such as the placozoans (Barber et al., 2009; Lek et al., 2012; Washington and Ward, 2006). In contrast, MCTPs were not identified using the structural criteria listed above. Unicellular organisms present C2 proteins that have from one to seven C2 domains, including PKCs, ferlins and tricalbins (Barber et al., 2009; Lek et al., 2010; Schulz and Creutz, 2004). Thus, it can be suggested that MCTPs emerged as an adaptation in metazoans, but we do not know yet what role they may play. To understand how the proteins function in nature and how they evolve, it is important to explore shared similarities from amino acid sequences in different species to identify paralogs and homologs (Bolker, 2014; Lek et al., 2010; Train et al., 2017). We also identified highly conserved regions (C2 domains and TMRs) and found that the sequences of the C2B and TMR1 domains have more amino acid substitutions compared to the C2A,C2C an TMR2.

X-ray diffraction analysis indicated that C2 domains are composed of eight-stranded β-sandwich with calcium-binding sites coordinated by aspartate residues (Rizo and Sudhof, 1998; Sutton et al., 1995). We analyzed the sequences of the three MCTP C2 domains to try to understand the structural array of these domains comparing them with the C2A and C2B domains of Synaptotagmin-1. The secondary structure is highly conserved in all sequences from selected organisms. Our results are similar to those reported by Shin et al. (2005) who showed that several organisms lack the calcium-binding aspartates in the C2B domain. Interestingly, we also identified that *C. elegans. H. contortus, X. laevis, X. tropicalis, M. musculus and H. sapiens* lack two calcium-binding sites in the C2B, this suggests that this domain does not bind calcium in all species. MCTPs have been described as a novel family of proteins that can bind calcium independently of phospholipid complexes (Shin et al., 2005); however, we identified that C2A and C2B have a lysine-rich domain (a polybasic cluster) or the β-groove motif previously identified in the synaptotagmin-1 C2B domain (**P21579.1**) (Corbalan-Garcia and Gómez-Fernández, 2014). Some studies have demonstrated that this motif is important for interactions with inositol polyphosphates (IP_4_, IP_5_ and IP_6_) (Kojima, 1995) and PIP_2_, which are directly implicated in exocytosis, vesicle fusion, membrane trafficking, cytoskeleton dynamics and membrane permeability, as well as in the transport of membrane channels (Paolo and Camilli, 2006). It seems that this motif acts as a signal for C2 domain proteins to target membranes. It also suggests that MCTPs could bind different phospholipids from those reported by Shin et al. (2005). In the structural model here presented at least two potential binding sites for PIP_2_ in the polybasic clusters are disclosed; this opens the possibility that upon PIP_2_ binding these domains bring closer the C2 domains with the TMR. This requires further experimental evidence. Meanwhile the question remains as to whether MCTPs bind or not phospholipids.

Regarding the TMRs, we identified two of these domains in all analyzed sequences, except one human isoform that possess one single TMR. This isoform is probably the result of an alternative splicing event in exon 17 in the *MCTP-2* gene. The functional difference between MCTPs with one or two TMRs is currently not well understood; both have three C2 domains, thus they have the same potential to bind calcium and may play similar and/or complementary roles in the cell.

Thus, this differences could be essential for distinguishing different classes of MCTPs, and it might be related with the specialized function of each C2 domain such as in ferlins and Syts (Barber et al., 2009; Washington and Ward, 2006).

## Conclusions and future perspectives

In summary, we described the distribution of MCTPs in different phyla, analyzed in detail highly conserved regions and described their secondary structure. Our studies show that these proteins are widespread in metazoans and, because of their selective location in the endoplasmic reticulum and endocytic/exocytic pathways, they may play important roles in processes such as vesicle fusion, exocytosis and membrane trafficking, such as those of other C2 domain proteins. Finally, our analysis discloses unique characteristics of MCTPs that allows speculation on their function in phospholipid and calcium signaling and suggests a working hypothesis to test their function.

## Acknowledgments

This work was supported by CONACYT A1-S-7659 to AMT. José Luis Téllez Arreola is a doctoral student from Programa de Doctorado en Ciencias Biomédicas, Universidad Nacional Autónoma de México (UNAM) and received fellowship 395834 from CONACYT and Fulbright-Garcia Robles (COMEXUS). Jessica González Norris kindly edited the manuscript.

